# Neuronal control of suppression, initiation and completion of egg deposition in *Drosophila melanogaster*

**DOI:** 10.1101/2021.08.23.457359

**Authors:** Cristina Oliveira-Ferreira, Miguel Gaspar, Maria Luísa Vasconcelos

**Affiliations:** Champalimaud Research, Champalimaud Center for the Unknown, Lisbon, 1400-038, Portugal

## Abstract

Egg-laying in *Drosophila* is the product of post-mating physiological and behavioural changes that culminate in a stereotyped sequence of actions. While egg-laying behaviour has been mostly used as a system to understand the neuronal basis of decision making in the context of site selection, it harbours a great potential as a paradigm to uncover how, once a site is selected, the appropriate motor circuits are organized and activated to deposit an egg. To study this programme, we first describe the different stages of the egg-laying programme and the specific actions associated with each stage. Using a combination of neuronal activation and silencing experiments we characterize the role of three distinct neuronal populations in the abdominal ganglion with different contributions to the egg deposition motor elements. Specifically, we identify a subset of glutamatergic neurons and a subset of cholinergic neurons that promote the initiation and completion of egg expulsion respectively, while a subset of GABAergic neurons suppresses egg-laying. This study provides insight into the organization of neuronal circuits underlying complex motor behaviour.

## Introduction

Motor execution of behavioural programmes must be tightly controlled so that the appropriate sequences of actions is executed once the internal state and environmental cues are considered. Egg-laying is executed by female fruit flies up to 80 times per day^1^. For each egg laid, the female follows an egg-laying motor programme described as comprising a search-like period followed by egg deposition and subsequent clean and rest^2^. Presumably, during the search period the fly evaluates the environment in order to find the best site to deposit the egg. For flies and the other oviparous animals when and where an egg is deposited has a profound impact on the survival of the offspring. A female decides an egg laying site based on substrate texture^3,4^, food availability^2,5,6^ protection from climate elements^7,8^, predator avoidance^9–16^, and disease avoidance^17–20^. The female will also evaluate information from other flies^21–25^. In recent years a number of mechanosensory^3,4^, chemical^2,5–7,9–13,17,18,21^ and visual^8,14–16,23^ cues that modulate egg-laying site selection have been identified, creating a broad view of the external cues guiding egg-laying decisions. Internal cues prompt the search period in the female^26^. Movement of the egg through the reproductive system is in part controlled by octopamine, which contracts the ovaries and relaxes the oviduct, and glutamate which contracts the oviduct^27–29^. Sensory neurons expressing the mechanosensory channel *Piezo* at the oviduct detect contraction/distention leading to a search for an egg-laying site^26^. While great progress has been made in understanding site selection and its neuronal underpinnings, very little is known regarding the consummatory component of egg-laying: egg deposition.

Here we addressed the neuronal basis of the egg deposition motor programme, one of the phases of egg-laying, at the lower level of motor control. We began with a description of the behavioural elements of the full egg-laying motor programme in wild type flies. Next, we searched lines that label neurons in the abdominal ganglion of the ventral nerve cord where potentially brain commands and reproductive system inputs are integrated to perform egg deposition. Activation and silencing experiments revealed a line labelling neurons that are crucial for egg deposition behaviour. We reasoned that neurons expressing different neurotransmitters likely play distinct roles in executing egg deposition. Using an intersectional approach, we showed that cholinergic neurons are sufficient for egg pushing and expulsion, while glutamatergic neurons contribute for the initial elements of egg deposition. In contrast, GABAergic neurons suppress egg-laying. This study shows for the first time the link between the activity of abdominal ganglion neurons and specific behavioural motor elements. This knowledge provides insight into the architecture of the egg deposition circuitry at the level of the lower motor control.

## Results

### Characterization of egg-laying behaviour

In order to understand how the execution of egg-laying motor programme is coordinated we analysed wild type egg-laying behaviour in detail. Fig. 1a shows a schematic representation of the behavioural setup. Single females were placed in an arena lined with 1% agarose on three walls. Videos were recorded for 45 minutes. We first analysed the temporal structure of egg expulsion (Fig. 1b). For the duration of the video, each female laid from 4 to 17 eggs, with a median of 11.5 (Fig. 1c). All eggs were laid on the agarose walls and 94% of eggs were buried in the agarose. The median inter-egg expulsion interval calculated for each female ranges from 2 to 3 minutes with the exception of one fly (fly#7) that showed a median of 5 minutes (Fig. 1d). These data indicate a low inter-individual variability when rearing conditions and environment are controlled. To obtain a detailed description of egg-laying behaviour, we analysed egg-laying motor elements which we associated with different egg-laying phases. Egg-laying behaviour involves 1) an exploration phase where the fly presumably searches for appropriate egg-laying sites, 2) the egg deposition phase in which a motor programme is activated to lay an egg, and 3) and rest phase. We chose to call the rest phase ‘abdominal contortions’ because we and others^30,31^ observed that this phase is accompanied by strong and prolonged abdominal contortions, as described below. Fig. 1e shows videoframes capturing each of the different egg-laying motor elements. Some behavioural denominations were based on other descriptions of egg-laying^30,31^. Egg deposition motor elements all progress in a sequence culminating in egg expulsion and ending with grooming of the terminalia. We show more than one videoframe for egg pushing as it has two different postures. Abdominal contortions, as mentioned earlier, are the single motor element of its phase. We defined exploration motor elements as elements where females probe the substrate either with the ovipositor or the proboscis without progressing in continuous sequence to egg expulsion. To understand the temporal sequence of the different behavioural elements we plotted the probability of each motor element around the moment of egg expulsion (Fig. 1f-i). During egg deposition phase the motor elements leading to egg expulsion are performed every time an egg is deposited (Fig. 1f). After an egg is expelled, the female curls the abdomen while walking away which can be followed by grooming terminalia (Fig. 1g). While ovipositor contact, burrowing, egg pushing and abdomen curling are fixed elements of egg deposition, grooming terminalia is optional. A few seconds after the egg is expelled, the probability of abdominal contortions rises (Fig. 1h). The timing of abdominal contortions is very variable though abdominal contortions generally increase a few seconds after egg expulsion and decrease in the minute leading up to egg expulsion. It is likely that abdominal contortions correspond to the moment of ovulation since the next phase, exploration, is activated by ovulation^26^. As abdominal contortions subside, the exploration motor programme initiates (Fig. 1i). Females contact the substrate by extending the proboscis and by bending the abdomen to reach the surface with the extruded ovipositor. A small peak of proboscis extension is also observed after egg expulsion indicating there may also be an association of this behaviour with the post-egg expulsion motor sequence. Part of these ovipositor contacts are accompanied by burrowing. These two behavioural elements are the same as those observed in the egg deposition programme except that they are not accompanied by egg pushing or followed by egg expulsion. An overview of exploration and egg deposition phases together with the respective motor elements is provided in Supplementary Video 1. Abdominal contortions phase is represented in Supplementary Video 2. We next analysed the transition probabilities between the different phases (Fig. 1j). This analysis revealed that egg deposition is always followed by abdominal contortions. Abdominal contortions are either followed by exploration or transitions directly to egg deposition with similar likelihood. Once the female initiates exploration she does not return to abdominal contortions without depositing an egg first. Similarly, once the female deposits an egg, she does not explore without going through abdominal contortions first. In our quantification of phase transitions, we did not consider proboscis extension, as it is not exclusive to the exploration phase nor to egg-laying behaviour. Still, the results suggest that exploration is optional in this setup. Here we described seven behavioural elements and associated the elements with each phase, which sets the stage to analyse the neuronal circuits of egg-laying.

**Fig. 1.**
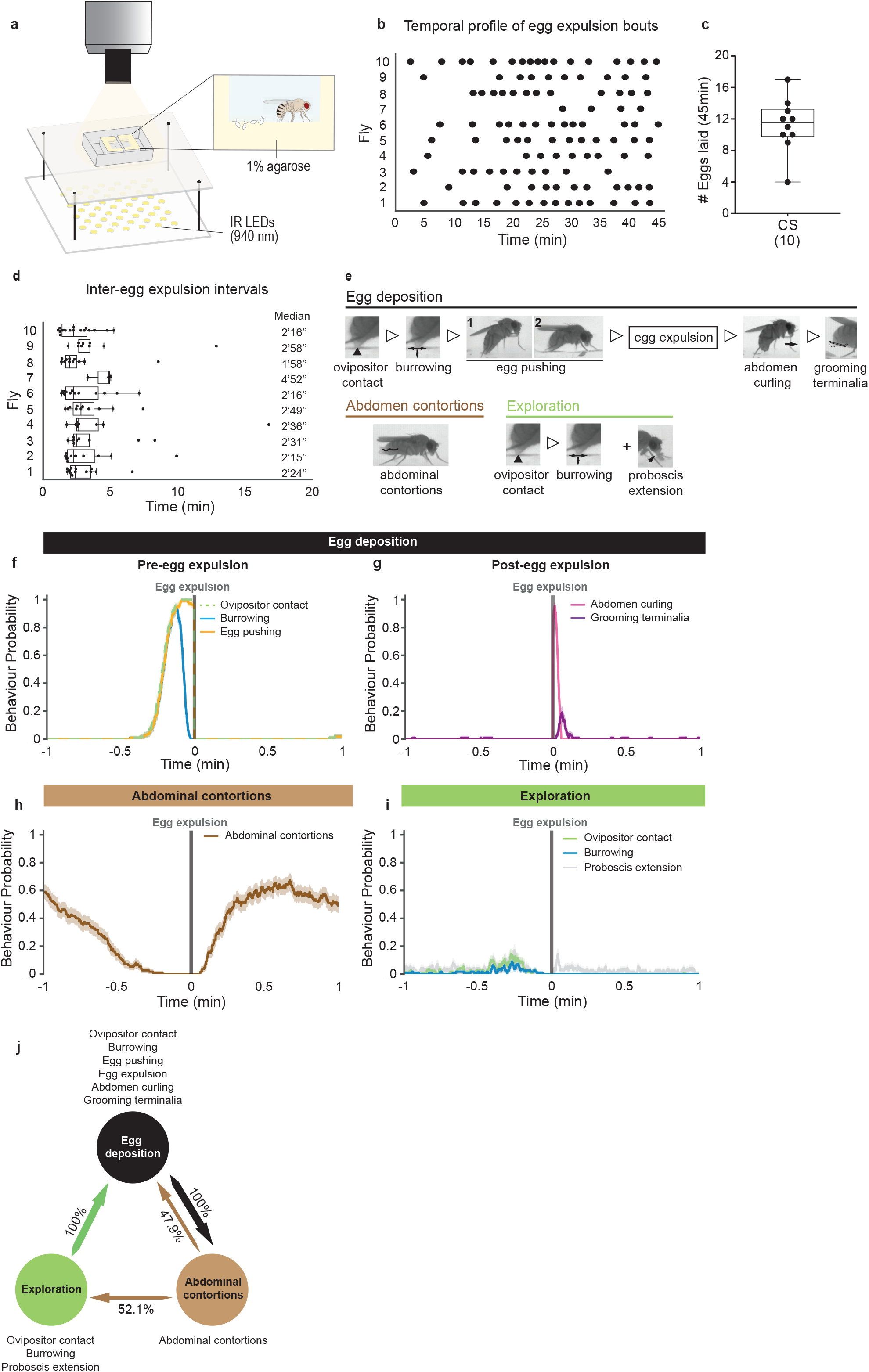
Egg-laying behaviour in wild type flies. **a** Schematic of setup and arenas used to record egg-laying behaviour. Each arena is composed by two chambers allowing to record two flies simultaneously. The egg-laying substrate used in this study was 1% agarose. Infrared (IR) LEDs were used for illumination and IR cameras were used to record fly behaviour. **b** Raster plot showing the temporal profile of egg expulsion bouts during a 45 min period. Each black dot marks the moment of egg expulsion. n = 10 flies. **c** Number of eggs laid by mated, egg-laying deprived, Canton S (CS) flies in 45 min videos. Median = 11.5; total number of flies = 10; total number of eggs = 112. **d** Time intervals between egg expulsion bouts per fly. Each dot represents the interval between consecutive egg deposition bouts. Median values of the distribution for each fly are shown on the right side. n = 10 flies. **e** Video snapshots illustrating the different egg-laying motor patterns analysed within each egg laying phase. Open triangles between snapshots denote sequential behaviours. Ovipositor contact is defined by abdomen bending accompanied by extrusion of the ovipositor and contact with underlying substrate (marked by the black arrowhead). It can culminate in egg expulsion or not. Burrowing is characterized by scratching surface eventually leading to digging the substrate with the ovipositor. Egg pushing is characterized by a rigid posture promoted by the contraction of the abdomen. Note that this behaviour can be displayed in two different postures: 1. erect body posture (fly head is elevated in relation to the abdomen, or 2. leaning body posture (fly leans towards the substrate). Abdomen curling is characterized by curling the abdomen followed by walking forward, either lifting the curled abdomen, or by dragging it through the substrate. Grooming terminalia is self-explanatory. Abdominal contortions are undulated abdominal movements (represented by the shape of the black line) accompanied by extrusion of the ovipositor. Proboscis extension is characterized by the proboscis being extended to contact the substrate (marked by the black arrow). **f**, **g**, **h** and **i** Probabilities of indicated female egg-laying behaviours during a 1-min time window around egg expulsion (time = 0, represented by the grey vertical line). Shaded area represents the standard error of the mean (SEM). n = 112 egg expulsions. **j** Transition probabilities of the egg-laying phases. Egg-laying behaviour comprises three distinct phases: egg deposition (black), exploration (green) and abdominal contortions (brown). Arrow thicknesses represent the degrees of transition likelihood. n = 10 flies.

### Activity of OvAbg neurons promotes egg deposition

Organs in the abdominal region of the flies connect to the central nervous system at the abdominal ganglion (Abg) of the ventral nerve cord (VNC). Since the reproductive organs are located in the abdomen and many egg-laying behavioural elements involve abdominal movement, this region is candidate to identify neurons involved in the execution of egg-laying behaviours. Therefore, we performed an activation screen of splitGal4 lines^32–34^ selected based on Abg expression and from this screen we selected a line, we will call OvAbg, for further investigation. Interestingly, the OvAbg line encompasses regulatory fragments of the sex determination gene, doublesex *(dsx)*, widely known to control sex specific reproductive behaviours in flies^34–38^. The OvAbg line anatomy reveals a group of neurons exclusively localized in the Abg (Fig. 2a). OvAbg neurons innervate other ganglia of the ventral nerve cord, the suboesophageal zone (SEZ) (Fig. 2b), and the reproductive system, specifically in the lateral oviducts and the distal uterus (Fig. 2c). Additionally, we found one OvAbg projection in the seventh abdominal segment of the body wall muscle (Supplementary Fig. 1a and b). To identify the polarity of OvAbg neurons we used synaptotagmin and Denmark labelling which revealed presynaptic terminals in the SEZ and mixed labelling of processes in the Abg and lateral oviducts (Supplementary Fig. 1c-k). To investigate the role of OvAbg neurons in egg-laying, we optogenetically activated them using CsChrimson^39^ with 6 stimuli of 10 seconds with a 20 second interval between stimuli (Fig. 2d). The neuronal activation always and only leads the mated female to assume an egg pushing posture (Fig. 2e, Supplementary Video 3). In contrast, the control females, which have the same genetic background as OvAbg but no regulatory element for splitAD^40^, never assume the egg pushing posture during light stimulation. Fig. 2f depicts the egg pushing posture of an OvAbg activated female and a wild type female during unmanipulated egg-laying for comparison. To address whether mating status affects the output of OvAbg neurons, we activated virgin OvAbg females. We observed that virgin females, like mated females, assume an egg pushing posture in all light ON periods (Fig. 2e). To test the contribution of the OvAbg projections in the brain to the activation phenotype, we activated headless females. We observed that similarly to intact OvAbg females, headless OvAbg females assume an egg pushing posture at every light stimulation (Fig. 2g and h), indicating that OvAbg brain projections do not contribute to this activation phenotype. Next, we quantified egg expulsion during the activation protocol. We observed that nearly half of the test flies expels an egg during the activation protocol whereas control females do not expel eggs (Fig. 2i). Most of the eggs are expelled during the first stimulation period (Fig. 2j). These results show that activity of OvAbg consistently leads to an egg pushing posture but does not always lead to egg expulsion. Given that OvAbg neurons control a female specific behaviour we asked whether these neurons are present in the male. Indeed, there are male OvAbg neurons albeit fewer and with dimorphic projections (Supplementary Fig. 1l-n). Activation of OvAbg neurons in males always triggers two motor elements associated with copulation, abdomen curling and aedeagus extrusion, suggesting an analogous role of OvAbg neurons in male reproduction (Supplementary Fig. 1o and p). To further investigate the role of OvAbg activity in egg-laying we performed silencing experiments using the inwardly rectifier potassium channel Kir2.1^41^. For egg-laying experiments females were paired with males on apple juice agar plates and monitored for two hours for copulation. The number of eggs laid by mated females over 24 hours was nearly abolished by OvAbg silencing (Fig. 2k). Notably, we observed that 88.7% test and 75% control flies mated showing that OvAbg neurons are not involved in female receptivity (Supplementary Fig. 1q). This result together with the observation that flies survive and appear healthy with constitutive silencing of OvAbg neurons indicates that they are specifically involved in egg-laying. To ascertain that egg-laying defect does not result from defect in egg production or ovulation, we dissected the reproductive system. We observed eggs jammed in the lateral oviducts in all test flies together with an excess of mature eggs in the ovaries (Fig. 2l and m), indicating that egg production and ovulation are not affected.

**Fig 2.**
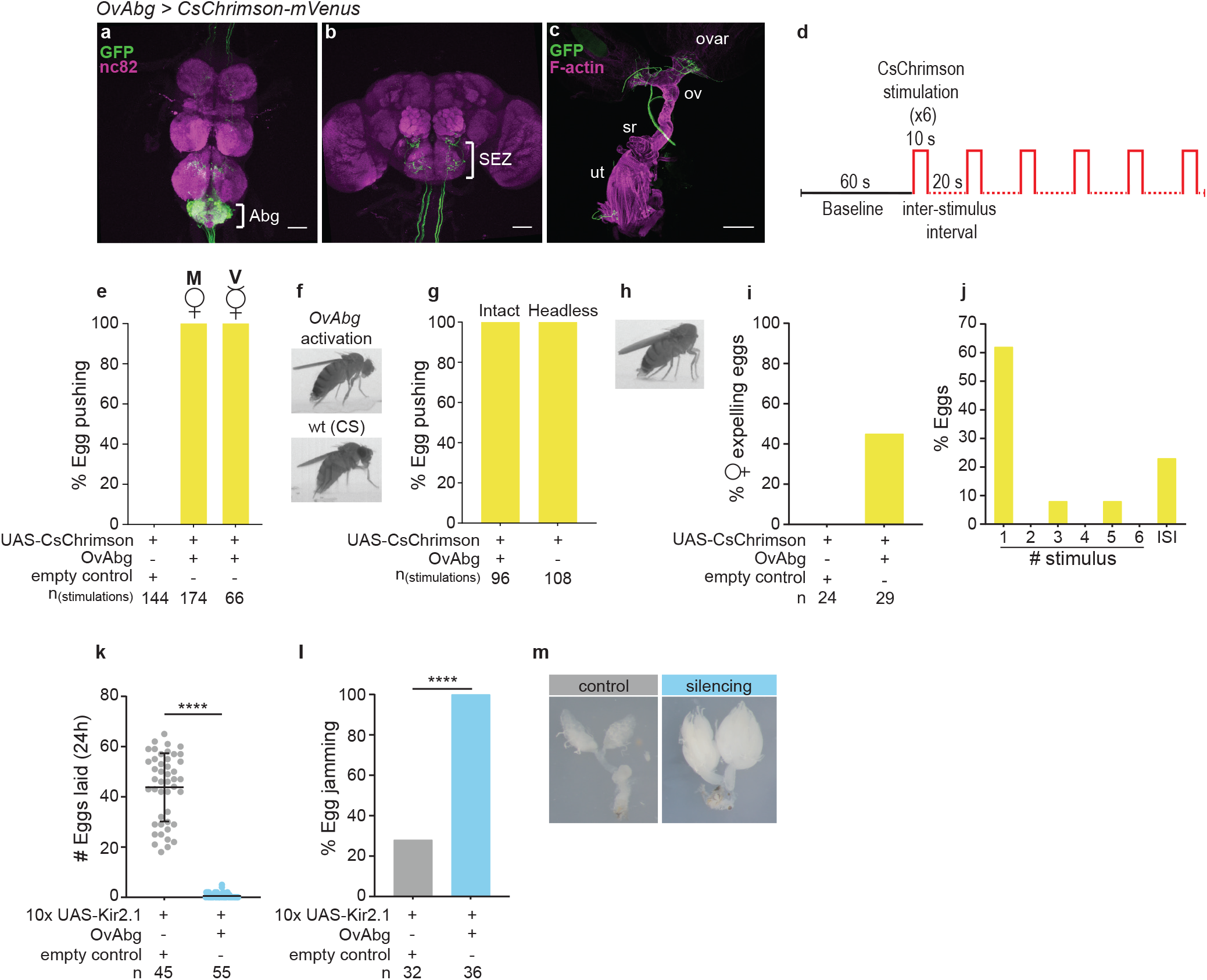
Activity of the abdominal ganglion neurons, OvAbg, promotes egg deposition. **a**, **b** and **c** Confocal images of female ventral nerve cord (VNC, (**a**)) brain (**b**) and reproductive system (**c**) of OvAbg neurons and corresponding innervations stained with anti-GFP (green) to reveal the OvAbg anatomy and nc82 for neuropil. Anti-F-actin was used in c) to visualize the muscle fibers. Abg: abdominal ganglion; SEZ: subesophageal zone; ovar: ovary; ov: oviducts; sr: seminal receptacle; ut: uterus. Anti-GFP is targeting the fluorescent protein Venus from OvAbg > CsChrimson-mVenus expressing flies. Scale bars a), b) 50 µm and c) 200 µm. **d** Neuronal CsChrimson activation protocol, which includes a baseline period (1 min) followed by 10 seconds activations repeated six times that are interspaced by 20 seconds intervals. **e** Percentage of stimulation events in which mated (M) and virgin (V) OvAbg flies displayed an egg pushing-like posture during photoactivation with CsChrimson. n = 144 (control) and 174 (OvAbg) stimulations. **f** Video snapshots (lateral view) of an OvAbg female (top) displaying an egg pushing-like posture in response to the stimulation with CsChrimson. A snapshot of a CS fly is also shown (bottom) for comparison of the egg pushing posture performed by wild-type (wt) flies during natural egg-laying behaviour. **g** Percentage of stimulation events in which headless OvAbg flies displayed an egg pushing posture during photoactivation with CsChrimson. n = 96 (control) and 108 (OvAbg) stimulations. **h** Video snapshot (lateral view) of a headless OvAbg female performing egg pushing-like posture in response to the stimulation with CsChrimson. **i** Percentage of OvAbg females laying eggs during photoactivation with CsChrimson. n = 24 (control) and 29 (OvAbg) females. **j** Percentage of eggs laid by OvAbg females during stimulation periods and inter-stimulus intervals (ISI). n = 13 eggs. **k** Number of eggs laid per female in the 24h after mating during nhibition of OvAbg neurons. n = 45 (control) and 55 (OvAbg) females. Mean ± s.d. is shown on top the scatter plot. Mann-Whitney test, ****p < 0.0001. **l** Percentage of females with eggs jammed in the lateral oviducts during inhibition of OvAbg neurons. n = 32 (control) and 36 (OvAbg) females. Fisheŕs exact test, ****p < 0.0001. **m** Image representing an OvAbg female reproductive system (right) with two eggs jammed (white arrows) in the lateral oviducts and a control (left) reproductive system with no eggs in the oviducts. Note that, besides the egg arresting, OvAbg silenced females also have enlarged ovaries containing more mature oocytes when compared with control ovaries.

In this series of experiments, we found a group of neurons that is part of the egg-laying motor circuits as they are directly involved in the local execution of egg pushing and egg expulsion and that are necessary for egg-laying. These Abg neurons provide a great entry point to address how different local circuits coordinate to execute egg deposition.

### Silencing OvAbg neurons disrupts all motor elements associated with egg-laying behaviour

We have shown that upon activation of OvAbg neurons a single egg deposition motor element - egg pushing - is induced and that OvAbg silenced females do not lay eggs. How do OvAbg silenced females behave? Do they perform all the behaviour elements with the exception of egg pushing or are other motor elements are affected? To answer these questions we used the anion channelrhodopsin GtACR1^42^ for acute optogenetic silencing of OvAbg neurons. We analysed 15 minutes of light stimulation as well as 10 minutes pre- and 5 minutes post-stimulation (Fig. 3a). Acute silencing of OvAbg neurons blocked egg-laying; the number of eggs laid during the stimulation was severely reduced and partially recovered in the post-stimulation period (Fig. 3b). Analysis of the behavioural elements showed that, with the exception of grooming, all egg deposition motor elements are abolished during stimulation and partially recovered post-stimulation (Fig. 3c-g). The expulsion of four eggs during the silencing period was done without using most of the egg deposition motor programme (Supplementary Video 4). The number of terminalia grooming bouts does not differ from control during silencing (Fig. 3g). However, the time the female spent grooming the terminalia is much larger in the test condition during silencing (Fig. 3h). Interestingly, both measures of grooming are reduced compared to control in the post-stimulation period suggesting a rebound effect on circuits modulating grooming behaviour. The data so far show a wide effect of silencing OvAbg neurons in all the egg deposition elements. We next analysed the other egg-laying phases. Abdominal contortions are reduced compared to control, both in number of bouts (Fig. 3i) and behaviour duration (Fig. 3j). Additionally, the intensity of the contortions and the extent of the ovipositor extrusion are reduced in test flies compared to controls, as exemplified in Fig. 3k. The behaviour elements of the exploration motor programme are reduced during silencing but fully recover post-stimulation (Fig. 3l-n) as do abdominal contortions, indicating that the regulation of these phases is simpler than that of the egg deposition programme where the inhibitory effects of GtACR stimulation persist.

**Fig. 3.**
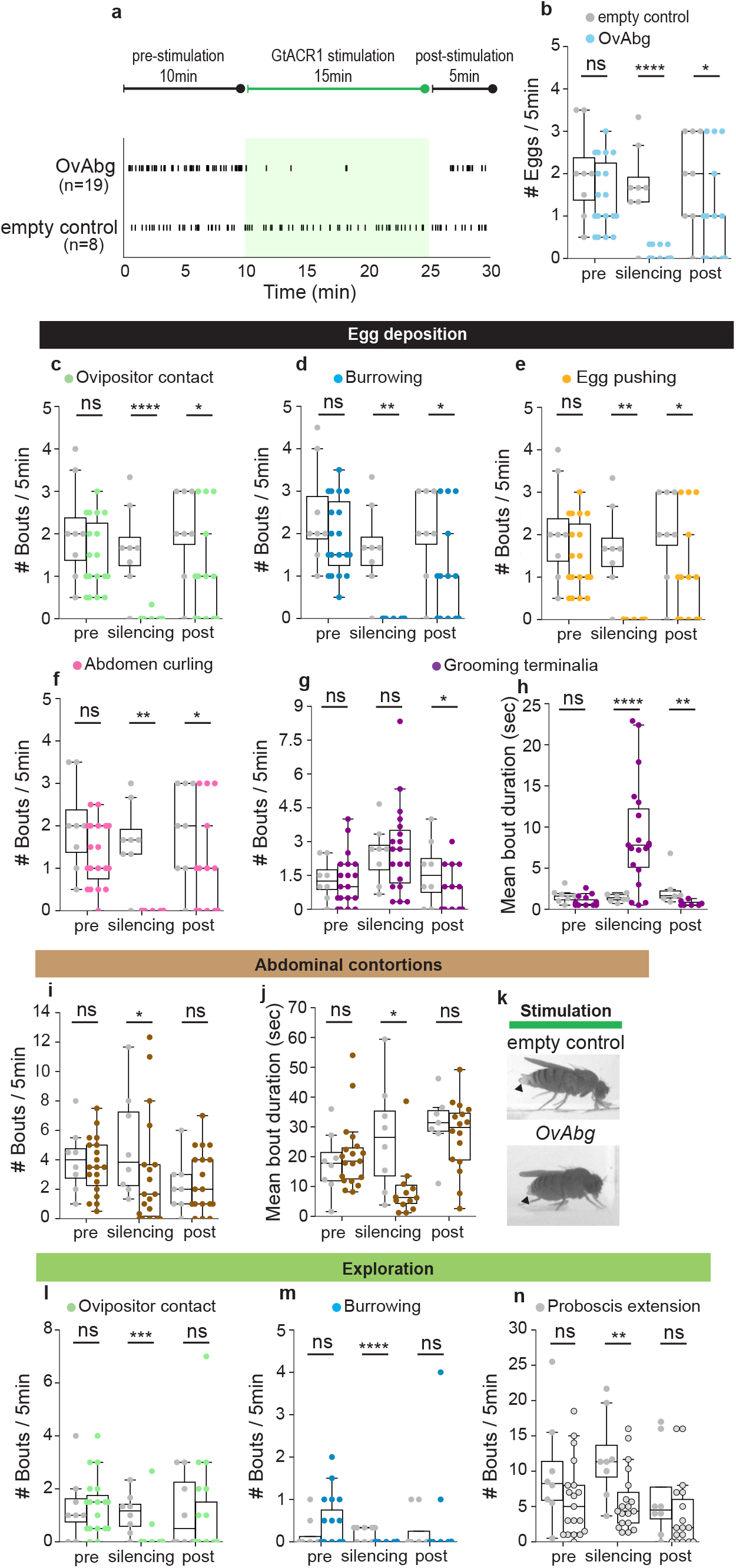
Silencing OvAbg neurons disrupts all motor elements associated with egg-laying behaviour. **a** (top) Neuronal GtACR1 silencing protocol scheme includes a pre-stimulation period (10 min), a silencing period (15 min) followed by a post-stimulation period (5 min). (bottom) Raster plot shows the egg expulsion events for OvAbg GtACR1 silenced and control females throughout the 30 minutes of the experimental protocol. Green shaded area represents the stimulation period. n = 19 (OvAbg) and 8 (control) flies. **b** Quantification of the number of eggs laid per fly by OvAbg and control females during 5 min periods. n = 19 (OvAbg) and 8 (control) flies. Pre: Mann-Whitney test, ns p ≥ 0.05; Silencing: Mann-Whitney test, ****p < 0.0001; Post: Mann-Whitney test, *p < 0.05. **c, d, e, f** and **g** Egg deposition phase-associated motor elements and corresponding quantification of the number of behaviour bouts during 5 min periods. n = 19 (OvAbg) and 8 (control) flies. Pre: Mann-Whitney test in **c**, **e** and t-test in **d, f, g** ns p-value ≥ 0.05; Silencing: Mann-Whitney test in **e**, **g**, ****p < 0.0001 and ns p ≥ 0.05 and t-test in **c**, **d**, **f**, **p < 0.01; Post: Mann-Whitney test in **c**, **d**, **e, f, g**, *p < 0.05. **h** Quantification of the mean duration of grooming terminalia bouts. n = 19 (OvAbg) and 8 (control) flies. Pre: Mann-Whitney test ns p-value ≥ 0.05; Silencing: t-test, ****p < 0.0001; Post: Mann-Whitney test **p < 0.01. **i** and **j** Quantification of the number of abdominal contortions bouts during 5 min periods and the corresponding bout mean duration. n = 19 (OvAbg) and 8 (control) flies. Pre: t-test in **i** and Mann-Whitney test in **j**, ns p ≥ 0.05; Silencing: Mann-Whitney test in **i** and **m** and **j** *p < 0.05; Post: Mann-Whitney test in **i** and t-test in **j**, ns p ≥ 0.05. **k** Video snapshots of test (bottom) and control (top) flies displaying abdominal contortions during the silencing period. Besides displaying shorter bouts of abdominal contortions, silenced flies also display less extended ovipositor extrusions during abdominal contortions in comparison with control flies that perform full ovipositor extrusions (arrowheads). **l**, **m** and **n** Exploration phase-associated motor elements and corresponding quantification of the number of behaviour bouts during 5 min periods. n = 19 (OvAbg) and 8 (control) flies. Pre: t test in **l** and Mann-Whitney test in **m** and **n**, ns p ≥ 0.05; Silencing: Mann-Whitney test in **l**, **m**, and **n**, **p < 0.01, ***p < 0.001, ****p < 0.0001; Post: Mann-Whitney test in **l**, **m**, and **n**, ns p ≥ 0.05.

Our findings show that silencing OvAbg neurons affects all phases of egg-laying behaviour. This dramatic result could reflect a direct involvement of OvAbg neurons in all egg-laying phases. Alternatively, they could reflect an arrest on the egg-laying cycle (Fig. 1j) induced by the loss of egg pushing behaviour and inability to complete egg deposition.

### Activity of GABAergic OvAbg neurons blocks egg-laying

To address how different neurons within OvAbg population contribute to the execution of egg-laying, we used an intersectional approach to obtain functional subgroups^43^. The GABAergic OvAbg (OvAbg/Gad1) neurons will be discussed here while cholinergic and glutamatergic OvAbg neurons will be discussed in the ensuing sections. OvAbg/Gad1 neurons (∼99 neurons, n=4 flies) are mostly local interneurons with sparse and faint projections to other VNC ganglia (Fig. 4a). No projections of OvAbg/Gad1 neurons were observed in the brain (Fig. 4b) or the reproductive system (Fig. 4c). Silencing OvAbg/Gad1 had no effect on the number of eggs laid in 24h (Fig. 4d). Therefore, if OvAbg/Gad1 neurons contribute to egg-laying they may do so by inhibiting egg-laying. To test this, we activated OvAbg/Gad1 neurons overnight (16h activation). We observed that activation of OvAbg/Gad1 abolishes egg-laying (Fig. 4e). Dissection of the ovaries at the end of the experiment revealed that the eggs are jammed at the lateral oviduct (Fig. 4f and g).

**Fig. 4.**
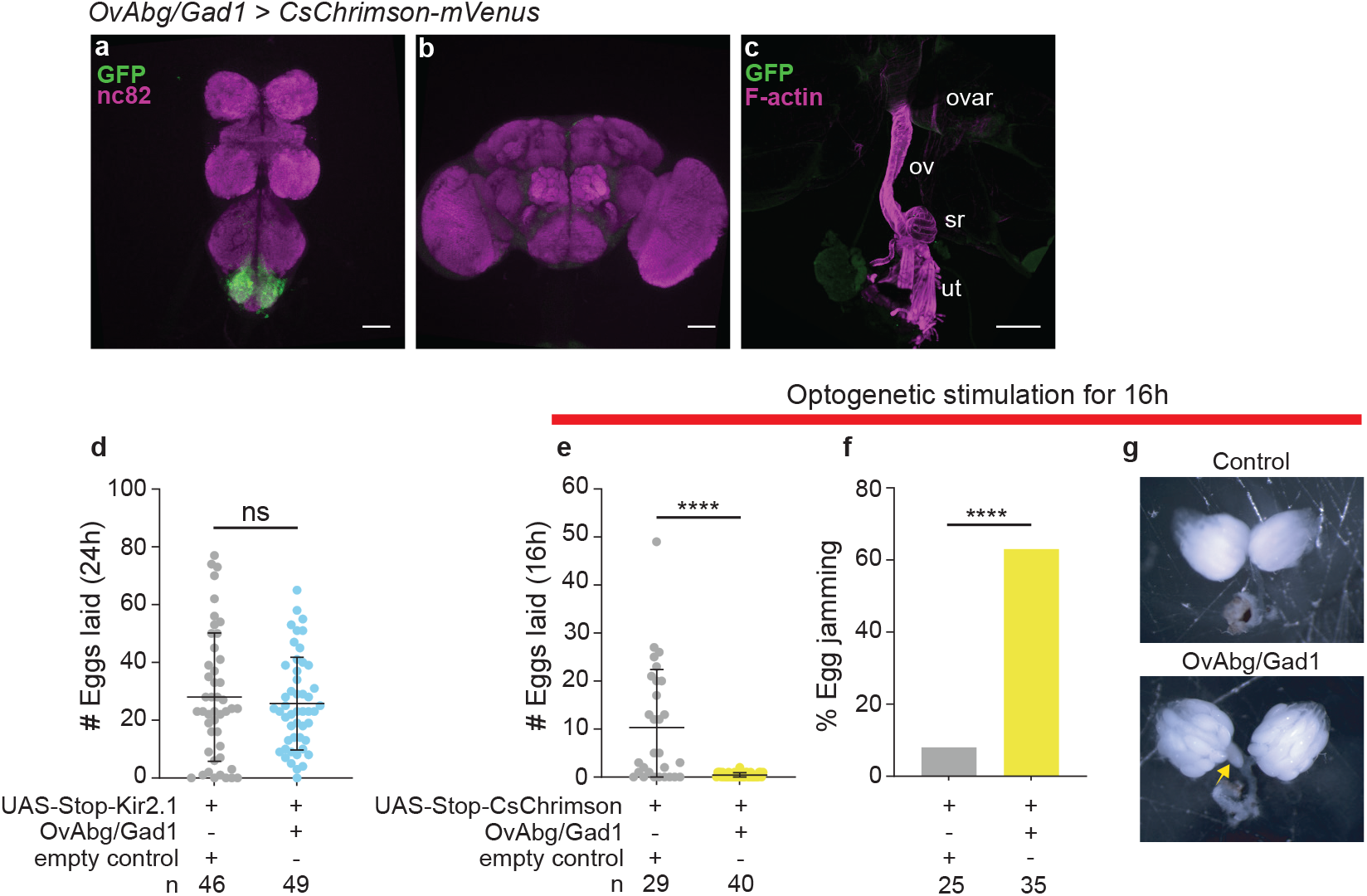
Activation of OvAbg GABAergic neurons is sufficient to block egg-laying. **a**, **b** and **c** Confocal images of female VNC (**a**), brain (**b**) and reproductive system (**c**) of OvAbg/Gad1 neurons and corresponding innervations stained with anti-GFP (green) to reveal the anatomy and nc82 for neuropil. Anti-F-actin was used in c) to visualize the muscle fibers. ovar: ovary; ov: oviducts; sr: seminal receptacle; ut: uterus. Anti-GFP is targeting the fluorescent protein Venus from OvAbg/Gad1-LexA > CsChrimson-mVenus expressing flies. Scale bars a), b) 50 µm and c) 200 µm. **d** Number of eggs laid per female in the 24h after mating during inhibition of OvAbg/Gad1 neurons. n = 46 (control) and 49 (OvAbg/Gad1) females. Mean ± s.d. is shown on top of the scatter plot. Mann-Whitney test, ns p ≥ 0.05. **e** Number of eggs laid per female during the 16h photoactivation with CsChrimson of OvAbg/Gad1 neurons. n = 29 (control) and 40 (OvAbg/Gad1) females. Mean ± s.d. is shown on top of the scatter plot. Mann-Whitney test, ****p < 0.0001. **f** Percentage of females with eggs jammed in the lateral oviducts after 16h photoactivation of OvAbg/Gad1 neurons. n = 25 (control) and 35 (OvAbg/Gad1) females. Fisheŕs exact test, ****p < 0.0001. **g** (bottom) Image representing a reproductive system of a OvAbg/Gad1 female after 16h photoactivation with one egg jammed (yellow arrow) in the lateral oviduct and a control (top) reproductive system with clear oviducts. Note that OvAbg/Gad1 activated females also have enlarged ovaries containing more mature oocytes when compared with control ovaries (similar to OvAbg silencing egg jamming phenotype, see figure 2l).

In summary, the results show that during egg-laying OvAbg/Gad1 neurons are silent and that activity in OvAbg/Gad1 neurons prevents egg-laying. This subset of OvAbg neurons contributes to opposing outcomes compared to the general line and thus have the potential to gate egg-laying execution by other neurons in the OvAbg population.

### Cholinergic OvAbg neurons are necessary and sufficient for egg deposition

Cholinergic neurons are a large fraction of OvAbg neurons (Fig. 5a) that include projections to the brain (Fig. 5b) and the reproductive system (Fig. 5c). Silencing cholinergic OvAbg (OvAbg/Cha) neurons leads to a severe reduction in the number of eggs laid (Fig. 5d) and a large fraction of the females display egg jamming in the lateral oviduct (Fig. 5e). These results show a very similar phenotype to that observed when silencing all OvAbg neurons (Fig. 2j and k). Likewise, activation of OvAbg/Cha neurons using the protocol shown in Fig. 2d leads to both virgin and mated females assuming an egg pushing posture each time the light is ON (Fig. 5f and g, Supplementary Video 5), as observed when of all OvAbg neurons are activated (Fig. 2d and e). Interestingly, quantification of females laying eggs during the stimulation protocol revealed that all OvAbg/Cha females laid one egg (Fig. 5h), in contrast to less than half of OvAbg females (Fig. 2h). Additionally, all eggs laid during the stimulation protocol by OvAbg/Cha females were laid during the first stimulus (Fig. 5i), whereas egg-laying timing by OvAbg females during the stimulation protocol was variable, with females laying eggs in the third and fifth stimulus as well as in the interstimulus intervals (Fig. 2i). The results show that, upon activation, if and when an egg is laid is variable for OvAbg, but not for OvAbg/Cha females. We also quantified, within the 10 second stimulation bout, when the females expelled the egg. We found a striking difference between OvAbg and OvAbg/Cha females (Fig. 5j), with OvAbg females taking a longer time to expel the egg. The increased variability regarding when the egg is expelled during the stimulation protocol and the increased latency to lay an egg upon light ON of the OvAbg females compared to the OvAbg/Cha females likely results from inhibition by the GABAergic neurons in the OvAbg population. Activation of OvAbg neurons encompasses simultaneous activation of inhibitory OvAbg/Gad together with egg-laying promoting OvAbg/Cha which results in conflicting information leading to variability and delay of the behavioural execution.

**Fig. 5.**
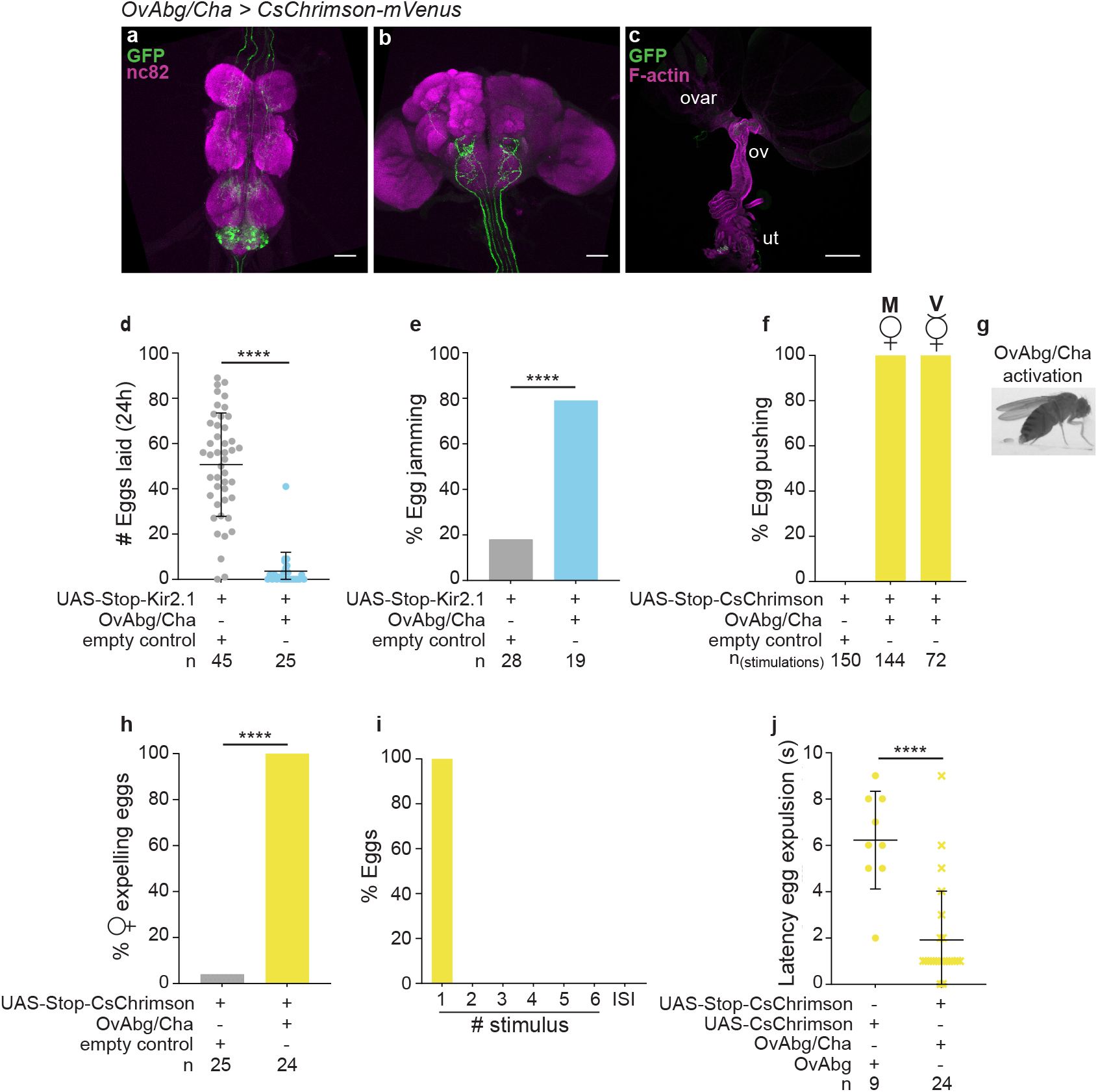
OvAbg Cholinergic neurons are involved in egg pushing and expulsion. **a**, **b** and **c** Confocal images of female VNC (**a**), brain (**b**) and reproductive system (**c**) showing OvAbg/Cha neurons and corresponding innervations stained with anti-GFP (green) to reveal the membranes and nc82 for synapses. Anti-F-actin was used in c) to visualize the muscle fibers. ovar: ovary; ov^1^: lateral oviducts; ov^2^: common oviducts; ut: uterus. Anti-GFP is targeting the fluorescent protein *Venus* from *OvAbg/Cha-LexA > CsChrimson-mVenus* expressing flies. Scale bars a), b) 50 µm and c) 200 µm. **d** Number of eggs laid per female in the 24h after mating during inhibition of OvAbg/Cha neurons. n = 45 (control) and 25 (OvAbg/Cha) females. Mean ± s.d. is shown on top of the scatter plot. Mann-Whitney test, ****p < 0.0001. **e** Percentage of females with eggs jammed in the lateral oviducts during inhibition of OvAbg/Cha neurons. n = 28 (control) and 19 (OvAbg/Cha) females. Fisheŕs exact test, ****p < 0.0001. **f** Percentage of stimulation events in which OvAbg/Cha flies displayed an egg pushing-like posture during photoactivation with CsChrimson. n = 150 (control) and 144 (OvAbg/Cha) stimulations. **g** Video snapshot (lateral view) of an OvAbg/Cha female displaying an egg pushing-like posture in response to the stimulation with CsChrimson. **h** Percentage of OvAbg/Cha females that lay eggs during photoactivation with CsChrimson. n = 24 (control) and 24 (OvAbg/Cha) females. Fisheŕs exact test, ****p < 0.0001. **i** Percentage of eggs laid by OvAbg/Cha females during stimulations and inter-stimulus intervals (ISI). n = 24 eggs. **j** Latency (seconds) to egg expulsion (period of time until the first egg-laying event after stimulus is ON)) of OvAbg and OvAbg/Cha photoactivated females. n = 9 (OvAbg) and 24 (OvAbg/Cha) females. Mean ± s.d. is shown on top of the scatter plot. Mann-Whitney test, ****p < 0.0001.

Overall, the results indicate that OvAbg/Cha neurons underlie the execution of egg pushing leading to egg expulsion. Unlike egg-laying behaviour of wild type flies, eggs expelled by optogenetically activated OvAbg/Cha females were never buried and were equally distributed between the acrylic and the agarose surfaces, highlighting the importance of other behavioural components for egg-laying site selection and egg burial.

### Glutamatergic OvAbg neurons contribute to egg deposition initiation

We found that glutamatergic OvAbg (OvAbg/VGlut neurons) is the smallest functional group composed by ∼10-18 OvAbg neurons (n=4 flies). We observed projections in the VNC, some unilateral and some bilateral, and occasionally we observed projections in the brain (Fig. 6a and b). This group of neurons also projects to the lateral oviducts and distal uterus (Fig. 6c). Optogenetic activation of OvAbg/VGlut neurons using the protocol depicted in Fig. 2d in mated females triggered ovipositor contact-related behaviours (Supplementary Video 6). We found that 55% of the stimulations lead to ovipositor contact behaviour (Fig. 6d and e, yellow box) while incomplete ovipositor contacts (ovipositor not touching the substrate) (Fig. 6e, pale yellow box) were identified in 21% of the stimulation periods (Fig. 6d). Additionally, in a small fraction of the stimulations (7%) flies displayed burrowing behaviour accompanying ovipositor contacts (Fig. 6d). Finally, we observed in 14% of stimulations flies either extruded the ovipositor with straight abdomen or bent the abdomen without extruding the ovipositor. Control flies did not display any of these behaviours during light ON (Fig. 6d). Interestingly, we found that virgin and mated OvAbg/VGlut females display different behavioural phenotypes upon activation. In virgins, 66% of the stimulations do not evoke any behaviour (Fig. 6d). Ovipositor extrusion or abdomen bending was observed in 18% of the stimulations. Ovipositor contact was identified in only 15% of the stimulations and burrowing behaviour displaying was residual (1% stimulations) (Fig. 6d). This result suggests that the behavioural output of OvAbg/VGlut neurons is modulated by the mating status. In the first section of this study, we characterized ovipositor contact and burrowing as motor elements displayed by pregnant females in the exploration and egg deposition phases. The phenotype of OvAbg/VGlut activation provides evidence for a role of OvAbg/VGlut in controlling these behaviours, which could be specific to one of those phases, or be implicated in both. To test this, we investigated in detail the egg-laying behaviour in OvAbg/VGlut females silenced with the potassium channel Kir2.1 since a GtACR tool that allows this type of genetic intersection is not available. We video-recorded silenced flies for 15 minutes and annotated their behaviour, as well as that of the control flies. Chronic silencing of OvAbg/VGlut neurons reduced, but did not abolish, the number of eggs laid (Fig. 6f), suggesting that this neuronal subset is not necessary for egg-laying in contrast to OvAbg/Cha group. Analysis of the behavioural elements shows that OvAbg/VGlut silencing affects egg deposition behaviours up to egg expulsion: ovipositor contact, burrowing and egg pushing (Fig. 6g, h and i). Silenced OvAbg/VGlut females displayed fewer bouts of each behaviour per egg, although we still observed manipulated flies that perform these behaviours at levels comparable to control. This reduction in the expression of motor elements that precede egg expulsion results in abnormal initiation of the egg deposition bout (Supplementary Video 7). Surprisingly, the post-egg expulsion behaviours - abdomen curling and grooming - were not reduced (Fig. 6j and k). In fact, grooming was slightly increased which may be associated with the deficient egg deposition (Fig. 6k). Indeed, the increased grooming in test females is specifically associated with the egg deposition events, rather than a generalised increase in grooming terminalia throughout the video (Supplementary Fig. S2). The contortions phase was not affected (Fig. 6l). Glutamate has been shown to be involved in oviduct contractions^28^, which presumably occur during the contortions phase. Our results indicate that there is a distinct group of glutamatergic neurons, not labelled by OvAbg, involved in oviduct contractions. The exploration phase, which shares two motor elements with the egg deposition phase (ovipositor contact and burrowing) and also includes proboscis extension was also not affected (Fig. 6m-o). Egg-laying in *Drosophila* culminates in egg expulsion subterraneously in the substrate. Since silencing OvAbg/VGlut neurons reduces pre-egg expulsion behaviours, we investigated if silenced females had less eggs buried. Indeed, by comparing the number of eggs dropped on the agarose surface (not buried) in relation to the total number of eggs laid, we observed that OvAbg/VGlut silenced females had less eggs buried in the agarose substrate (Fig. 6p) in comparison to control flies.

**Fig. 6.**
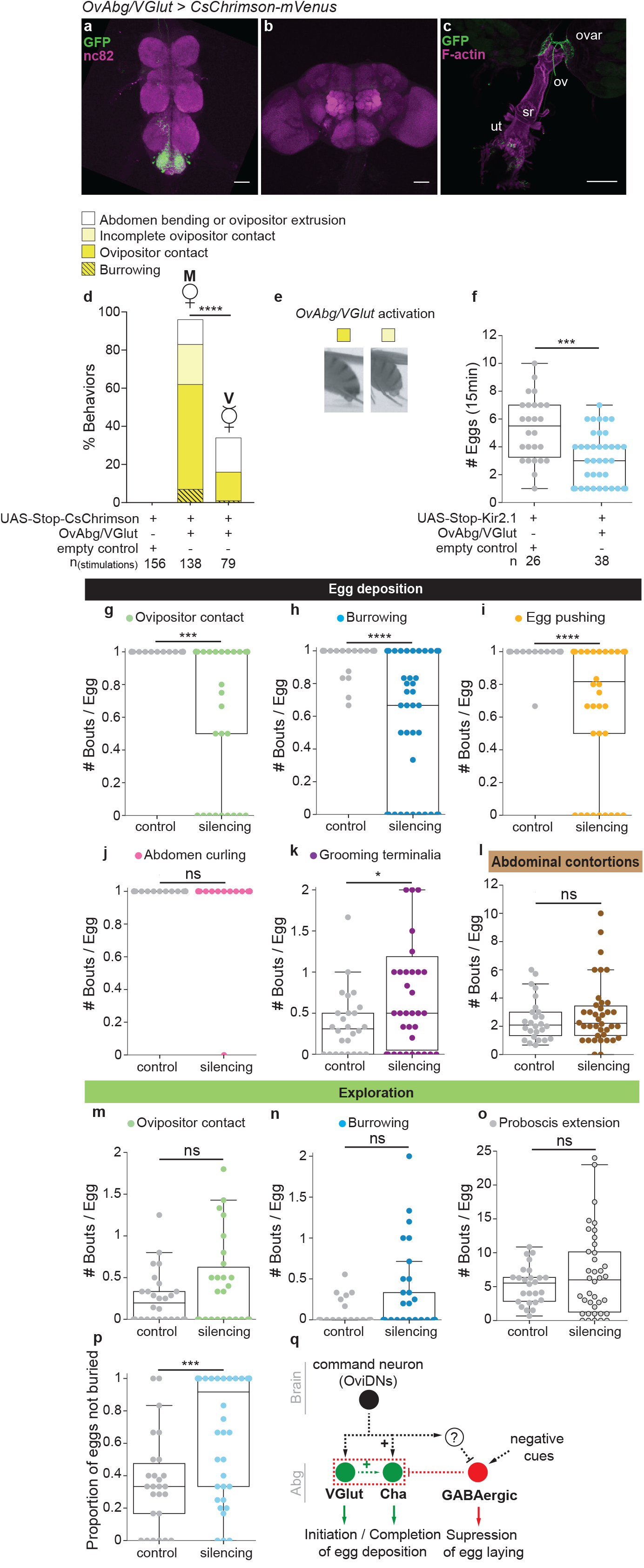
OvAbg Glutamatergic neurons contribute to egg deposition initiation. **a**, **b** and **c** Confocal images of female VNC (**a**), brain (**b**) and reproductive system (**c**) showing OvAbg/VGlut neurons and corresponding innervations stained with anti-GFP (green) to reveal the anatomy and nc82 for synapses. Anti-F-actin was used in c) to visualize the muscle fibers. ovar: ovary; ov: oviducts; sr: seminal receptacle; ut: uterus. Anti-GFP is targeting the fluorescent protein *Venus* from *OvAbg/VGlut-LexA > CsChrimson-mVenus* expressing flies. Scale bars a), b) 50 µm and c) 200 µm. **d** Percentage of stimulation events/periods in which OvAbg/VGlut mated (M) and virgin (V) flies displayed ovipositor contact, incomplete ovipositor contact, burrowing, abdomen bending or ovipositor extrusion behaviours during photoactivation with CsChrimson. n = 156 (control mated), 138 (OvAbg/VGlut mated) and 79 (OvAbg/VGlut virgin) stimulations analysed. Fisheŕs exact test, ****p < 0.0001. **e** Video snapshots (lateral view) of OvAbg/VGlut females displaying ovipositor contact and incomplete ovipositor contact behaviours in response to the stimulation with CsChrimson. **f** Number of eggs laid per female while silencing OvAbg/VGlut neurons for a period of 15 min. n = 26 (control) and 38 (OvAbg/VGlut) females. Mann-Whitney test, ***p < 0.001. **g**, **h**, **i**, **j** and **k** Egg deposition-associated motor elements and corresponding quantification of the number of behaviour bouts normalized for the number of eggs laid per female. n = 26 (control) and 38 (OvAbg/VGlut) flies. Mann-Whitney test in **g**, **h**, **i**, **j** and **k**, ****p < 0.0001, ***p < 0.001, *p < 0.05, ns p ≥ 0.05. **l** Quantification of the number of abdominal contortions bouts normalized for the number of eggs laid per female. n = 26 (control) and 38 (OvAbg/VGlut) flies. Mann-Whitney test, ns p ≥ 0.05. **m**, **n** and **o** Exploration phase-associated motor elements and corresponding quantification of the number of behaviour bouts normalized for the number of eggs laid per female. n = 26 (control) and 38 (OvAbg/VGlut) flies. Mann-Whitney test in **m**, **n** and **o**, ns p ≥ 0.05. **p** Proportion of eggs not buried per fly. n = 26 (control) and 38 (OvAbg/VGlut) flies. Mann-Whitney test, ***p < 0.001. **q** Proposed model of how glutamatergic, cholinergic and GABAergic OvAbg neurons may function in a local circuit to control and execute egg deposition behaviour. Solid arrows are based on our data and dashed arrows represent hypothetical connections.

In summary, the activation and silencing results strongly suggest that glutamatergic OvAbg neurons are involved in the initiation of the egg deposition motor sequence. Although OvAbg/VGlut neurons are not necessary for egg-laying, egg-laying is less efficient in OvAbg/VGlut silenced flies due to deficient egg deposition initiation. Furthermore, OvAbg/VGlut neurons play a role in progeny survival as silencing OvAbg/VGlut leads to fewer eggs buried in the substrate and, therefore, exposed to predation and adverse climate conditions.

## Discussion

Egg-laying is an excellent model for exploring the architecture of lower motor control as it is a complex behaviour constituted by different phases, each with a specific behavioural repertoire. In our observations, complementary to Yang et al^2^, egg-laying of wild type flies is structured in three phases - egg deposition, abdominal contortions and exploration. We showed that the egg deposition phase is always followed by contortions, which may be followed by a new egg deposition event or by an exploration period. This detailed description of the behaviour provided a framework to interrogate the underlying neuronal substrates. We identified three populations of OvAbg neurons in the abdominal ganglion with diverse contributions to the egg deposition phase. All three populations innervate similar areas in the abdominal ganglion, allowing potential connections between the three populations as part of an egg deposition circuit. Cholinergic neurons are involved in egg pushing leading to egg expulsion. Glutamatergic neurons are involved in the initiation of egg deposition. Interestingly, the initial motor elements of egg deposition that are shared with exploration-ovipositor contact and burrowing-were not affected in the exploration phase, suggesting different control mechanisms for the same motor elements during different phases. Our findings indicate that glutamatergic and cholinergic populations collaborate in the execution of egg deposition (Fig. 6q). In contrast, the GABAergic population suppresses egg-laying possibly acting on the glutamatergic and cholinergic populations (Fig. 6q). Our results indicate that in the conditions tested, the GABAergic population is silent during egg-laying as the number of eggs laid was not affected with silencing this population. Local suppression of egg-laying may be required when negative egg-laying cues arise, which could be mediated by an increase in activity in the GABAergic population.

The abdominal ganglion egg deposition circuit is poised to receive commands from the brain. The brain is thought to be involved in adaptive behaviour based on information collected from the environment and internal state. Thus, egg-laying site selection is processed in the brain which communicates with the abdominal ganglion for the execution of egg deposition. Indeed, many sensory inputs to the brain have been implicated in egg-laying site selection^2–18,21^. Brain neurons that may integrate these inputs and mating status information to make a decision are OviIN, OviEN and aDN^25,44^. An egg-laying decision is communicated to the abdominal ganglion through the descending neurons OviDNa and b^44^. We propose that cholinergic and glutamatergic OvAbg neurons receive the command from OviDNs to execute the motor actions of egg deposition (Fig. 6q).

Egg-laying is essentially performed by mated females though virgin females may deposit unfertilized eggs residually. Activation of OvAbg/VGlut neurons leads to fewer events of egg deposition initiation in virgin females when compared to mated females thus suggesting that mating status modulation of egg-laying is occurring locally at the abdominal ganglion in addition to the modulation in the brain^44–46^. Local modulation may result from direct octopaminergic modulation^47^ or in downstream targets.

In our work on egg-laying behaviour of wild type females, the exploration phase is defined by the ovipositor contact with the substrate, burrowing and the proboscis extension. In our assays, we see that the exploration phase is not obligatory. Two factors may explain this observation: 1) there may be exploration that does not include the motor elements we considered and rather flies use the mechanosensory and chemosensory information of the legs; 2) we use very small arenas with a restricted space for exploration and without complex sensory cues. This feature may allow a quick spatial and sensory recognition of the environment making exploration less frequent. Future work on the features of exploration and site selection should use more complex arenas and environments and consider kinematic analysis.

Our findings provide a detailed description of egg-laying. We described the different motor elements, their participation in different egg-laying phases and how flies transition from one phase to the next. This description facilitates the goal of linking a complex behaviour- egg-laying- with its neuronal underpinnings. Using our framework, we present insights into the logic of local egg deposition circuits. This work serves as a stepping stone to dissect ascending and descending communication with the brain, to extend neuronal dissection and connectivity of local egg-laying populations and address mechanisms of local mating status modulation of egg-laying.

## Material and Methods

### Fly stocks and husbandry

See Supplementary Table 1 and 2 for genotypes of *Drosophila* used in this study. Fruit flies *D. melanogaster* were raised in standard cornmeal-agar medium, using Vienna food recipe (In 1 Liter of water: 80 g molasses-barley malt, 22 g beet syrup, 80 g corn flour, 18 g granulated yeast, 10 g soy flour, 8 g agar-agar, 8 mL propionic acid,12 mL 15% nipagin, 35 mL Bavistin), at 25°C and 70% relative humidity in a 12 h dark:12 h light cycle. Detailed information on fly housing and age for each experiment are indicated in the relevant section.

### Immunohistochemistry

Adult brains, VNCs, reproductive systems and abdomen cuticles were dissected in cold phosphate-buffered saline (PBS) and immediately transferred to cold PFA 4% in PBL (PBS with 0.12 M Lysine) and fixed for 30 min at RT, washed three times for 5 min in PBT (PBS with 0.5% Triton X-100) and blocked for 30 min at RT in 10% normal goat serum in PBT (Sigma, cat# G9023). Samples were incubated with the primary antibodies in blocking solution, for 72 h at 4 °C. The following primary antibodies were used: rabbit anti-GFP 1:1000 (Molecular Probes, cat#A11122), chicken anti-GFP 1:1000 (abcam, ab13970), mouse anti-nc82 1:10 (Developmental Studies Hybridoma Bank, cat# AB_2314866), rabbit anti-DsRed 1:1000 (Takara, cat# 632496). Samples were washed three times for 5 min in PBT and incubated in Alexa Fluor 488 or 594 secondary antibodies 1:500 (Invitrogen) for 72 h at 4°C. To counterstain the female reproductive system, Alexa 594-conjugated phalloidin (Molecular Probes, cat# A12381) was used. Samples were washed three times for 5 min in PBT and mounted in VectaShield medium (Vector Laboratories, cat#H-1000). Images were acquired on a Zeiss LSM 710 confocal microscope using a 25X immersion objective (Zeiss) for the brains/VNCs and a 10X objective (Zeiss) for the reproductive systems and abdomen cuticles. After acquisition, colour levels were adjusted using Fiji^48^ for optimal display.

### Preparation of flies to be assayed

Low fly density crosses (10-15 virgin females x 5 males per bottle) were used in all experiments for rearing flies with the appropriate genotype. In order to maximize the occurrence of egg deposition events during behaviour experiments, we followed the egg-laying deprivation protocol described by Yang et al^49^ in all experiments, except in the 24h egg-laying assays. Briefly, groups of 5-7 virgin females per vial of the appropriate genotypes and 2-3 Canton S males (for mating) were collected into normal food vials with the exception of optogenetic experiments in which flies were housed in normal food containing all-trans-Retinal (Sigma, R2500) (all-trans-Retinal concentrations used: 0.2 mM for CsChrimson activation and 0.4 mM for GtACR1 silencing). In contrast with the original protocol^49^, yeast paste was not added to the food. Flies were left in the vials for 4 to 7 days at 25°C and 70% relative humidity. Behavioural assays were performed within that 4–7 days’ time window.

### Behavioural assays

#### 24h egg-laying assay

Single virgin females were gently aspirated and transferred to 35 mm Petri dishes of 10 mm of height (Thermo Fisher Scientific) coated with apple agar (750 mL water, 250 mL apple juice, 19,5 g agar, 20 g sugar, 10 mL 10% nipagin) and incubated with a naive Canton S male for 2 h under constant observation to check for mating occurrence. Plates where mating did not happen were discarded. Flies were kept in the plate for 24 h before eggs were counted. After egg counting, the femalés reproductive system was dissected to measure egg jamming.

#### Egg-laying arena and substrate

Custom made small rectangular-shaped arenas with 2 chambers were designed to allow recording of 2 flies simultaneously. Each chamber measures 1.8 (H) x 1.2 (L) x 0.3 (D) cm. During behavioural assays, the chambers were partially filled with the egg-laying substrate, which in this study was always 1% agarose (SeaKam® LE Agarose, cat# 50004) diluted in distilled water (Milli-Q® Water Purification Systems Merk). Flies had a free walking space of 1.5 x 0.7 x 0.3 cm in the chamber for egg-laying. The base and the lid of the arena were made of white opaque and transparent acrylic, respectively.

#### Detailed behaviour

To analyse the motor elements associated with egg-laying behaviour, females were collected soon after eclosion and housed in groups following the egg-laying deprivation protocol described above. Aged 4-7 days females were tested. Flies were gently aspirated into the egg-laying arena and behaviour was recorded at 20 frames per second during 45 min for Canton S (Fig. 1) and during 15 min for OvAbg/VGlut silenced females (Fig. 6f-p). The same fly handling procedure was performed for optogenetic experiments in which egg-laying motor elements were analysed (for more detailed information, see the optogenetic stimulation section).

#### Optogenetic stimulation

For all experiments using CsChrimson, except in the 16h egg-laying assay (Fig. 4e-g), the stimulation protocol included a 1 min baseline period followed by 6 repetitions of 10 s red-light stimuli with a power of 4.40 mW/cm^2^ and 20 s interval between stimuli. Fly behaviour was recorded at 20 frames per second, except in the OvAbg line activation experiments (Fig. 2e-j) in which we used 15 frames per second. In the OvAbg headless females’ photoactivation (Fig. 2g), the head was gently cut using dissection forceps (Dumont #55 Forceps, 11295-51) under CO_2_ anaesthesia. Flies were transferred to the egg-laying arena and allowed to recover from this procedure for 5-10 min before photoactivation. In the 16h photoactivation egg-laying assay (Fig. 4e-g), mated females were transferred to the apple agar plates (described in the 24 h egg-laying assay section). The stimulation protocol included constant red-light with a power ranging 4.19-4.85 mW/cm^2^ during 16 h. At the end of this period, the eggs were counted and the reproductive system was dissected to measure egg jamming. For the silencing experiment using GtACR1 (Fig. 3), the stimulation protocol included a pre-stimulation period that lasted for 10 min, followed by constant green-light stimulation with a power of 5-6.23 mW/cm^2^ during 15 min and a post-stimulation period of 5 min. Videos were recorded at 20 frames per second.

#### Image capture

Flies were filmed using a camera mounted above the arena (Teledyne Flir Flea3 FL3-U3-13S2M equipped with a 16mm fixed focal length lens (Edmund Optics)) and with a resolution of 1328 x 1048 pixels. For illumination, an infrared light using a 940 nm LED strip (SOLAROX) and a Hoya 49 nm R72 infrared filter (to reduce interference of visible light) were used. An electronic HARP LED array interface v1.3 and an HARP LED array v2.0 developed by the Champalimaud Foundation Hardware Platform were used to evoke a high-powered 610nm (CsChrimson activation) and 527nm (GtACR1 silencing) light. Bonsai software^50^ was used to trigger the optogenetics stimuli and to acquire the videos as avi files.

### Quantification and statistical analysis

#### Data processing

After movies were acquired, the in-house developed software Python VideoAnnotator was used to manually annotate the time and duration of all egg-laying motor elements under analysis. Annotations were done for the total duration of the video in all experiments.

#### Quantification of behaviours

Data and statistical analysis were performed using custom Python scripts in Fig. 1b-j, Fig. 3 and Fig. 6f-p. All the other analysis were performed using Prism9 (GraphPad Software, La Jolla, CA).

The inter-egg expulsion intervals (Fig. 1d) were calculated as

> Inter-egg expulsion intervals = first frame of egg expulsion bout - last frame of previous egg expulsion bout (per fly)

The number of eggs per 5 minutes (Fig. 3b) was calculated as

> # Eggs / 5 min = (sum # egg expulsions) / 5 minutes (per fly)

The number of behaviour bouts per 5 minutes (Fig. 3c-g, I, l-n) was calculated as

> # Bouts / 5 min = (sum # behaviour frames) / 5 minutes (per fly)

The number of behaviour bouts per egg (Fig. 6g-o) was calculated as

> # Bouts / Egg = (sum # behaviour frames) / total # egg expulsions (per fly)

The mean bout duration (Fig. 3h and j) was calculated as

> mean bout duration = # behaviour frame duration (s) / total # behaviour bouts (per fly)

Where the behaviour frame duration is given by the last frame subtracted to the first frame of each behaviour bout converted to seconds.

The proportion of eggs not buried (Fig. 6p) was calculated as

> Proportion of eggs not buried = (sum # eggs not buried) / total # eggs (per fly) Egg expulsion bouts in which it was not possible to determine whether the egg is buried or not were excluded from this analysis.

The percentage of behaviour displayed by flies during photoactivation (Fig. 2e-g, Fig. 5f) was calculated as

> % Behaviour = (sum # stimuli with behaviour) / total # stimuli

The percentage of females laying eggs (Fig. 2i and Fig 5h) was calculated as

> % Females laying eggs = (sum # females that laid eggs) / total # of females

The percentage of eggs laid during the photoactivation protocol (Fig. 2j and Fig. 5i) was calculated as

> % Eggs = (sum # eggs laid on each stimulus or ISI) / total # eggs

The latency for egg expulsion (Fig. 5j) was calculated as

> Latency egg expulsion = first frame corresponding to egg expulsion event – first frame corresponding to the stimulus initiation (per fly)

The product was converted to seconds.

The percentage of eggs jammed (Fig. 2l, Fig. 4f and Fig. 5e) was calculated as

> % Eggs jammed = (sum # reproductive systems with eggs in the oviducts) / total # reproductive systems

To investigate the egg-laying phases transition probability (Fig.1j), we first choose an egg-laying motor element to represent each phase: egg expulsion represented the egg deposition phase, abdominal contortions represented the abdominal contortions phase and ovipositor contacts that do not progress to egg expulsion represented the exploration phase. Proboscis extension and burrowing behaviours were not considered for the exploration phase transition analysis because: 1) proboscis extension displays a continuous occurrence during egg-laying behaviour (see Fig. 1i); 2) proboscis extension behaviour is not exclusive to egg-laying, being also displayed in other behavioural contexts (ex: feeding); 3) burrowing behaviour is displayed along with ovipositor contact but with lower frequency (see Fig. 1i). We calculated for the total video duration (45 min), the likelihood of all transitions between phases by using a first order Markov Chain analysis^51^. Phases transitions were identified as changes in the selected behavioural patterns for each phase.

To calculate the probability of the egg-laying behaviours around egg expulsion (Fig. 1f-i and Supplementary Fig. 2a and b) we aligned all the egg expulsion events of all flies and we counted how many behaviours were occurring in each of the 1200 frames preceding and following the end of egg expulsion. We then normalized the counts over the total number of egg expulsions. To represent the egg expulsion event, the last frame for each event was selected. We excluded from the analysis egg expulsion events that are less than 1200 frames from the start or end of the video.

#### Statistical analysis

Boxplot in Fig. 1c indicates the median flanked by the 25th and 75th percentiles (box) and whiskers showing the maximum and minimum values. In all the other boxplots the whiskers represent the 5th and 95th percentiles. Error bars when shown are mean ± standard deviation (SD). Prior to statistical testing, normality across all individual experiments was verified using the Shapiro’s Test, D’Agostino’s Test, Anderson-Darling test and Kolmogorov-Smirnov in Prism9 analysis. Only Shapirós and D’Agostinós tests were used in Python analysis. To assess the data variance homogeneity, Levenés and Bartlett’s tests were used for non-normally and normally distributed data, respectively. In all experiments we performed pairwise comparisons. Mann-Whitney test was used in non-normally distributed data. t-test was used in normally distributed data and the t-test with Welch’s correction was applied when data had unequal variances. Fisheŕs exact test was used in contingency tables.

The P value is provided in comparison with the control and indicated as * for p < 0.05, ** for p < 0.01, *** for p < 0.001, **** for p < 0.0001, and ‘ns’ for non-significant (p ≥ 0.05).

## Supporting information

Sup info Oliveira-Ferreira et al

## Acknowledgements

We thank the Champalimaud platforms for support during the development of this project. The fly facility for dissections, immunostainings, and fly food. The molecular and transgenic tools platform for screening recombinant flies. The scientific hardware platform for designing the HARP LED boards. The scientific software platform for developing Python VideoAnnotator. The ABBE platform for support with imaging. We thank Sophie Dias for support with fly husbandry. We thank Artur Silva and Matheus Farias for help with Bonsai software. Alexandre Azinheira and Tiago Coelho for editing the supplementary videos. We thank to Susana Lima, Ricardo Neto-Silva, Eugenia Chiappe, Michael Orger and members of Vasconcelos lab for feedback on the manuscript. This work was supported by Fundação Champalimaud and Portuguese national funds, through FCT—Fundação para a Ciência e a Tecnologia—in the context of the projects UIDB/04443/2020, Congento, LISBOA-01-0145-FEDER-02270, PPBI—LISBOA-01-0145-FEDER-022122 and BioData.pt— LISBOA-01-0145-FEDER-022231. C.O.F. was supported by FCT doctoral fellowship under the International Neuroscience Doctoral Programme (SFRH/BD/52448 / 2013).

## Author contributions

C.O.F. and M.L.V. conceived and designed the project. All experiments were performed and analysed by C.O.F., with the exception of Figures 1, 3 and 6 where M.G. contributed with python analysis. M.L.V. provided guidance and wrote the manuscript together with C.O.F.. M.G. reviewed the manuscript.

