## Supplementary material for "Neuronal control of suppression, initiation and completion of egg deposition in *Drosophila melanogaster*": Sup info Oliveira-Ferreira et al

Supplementary Fig.1

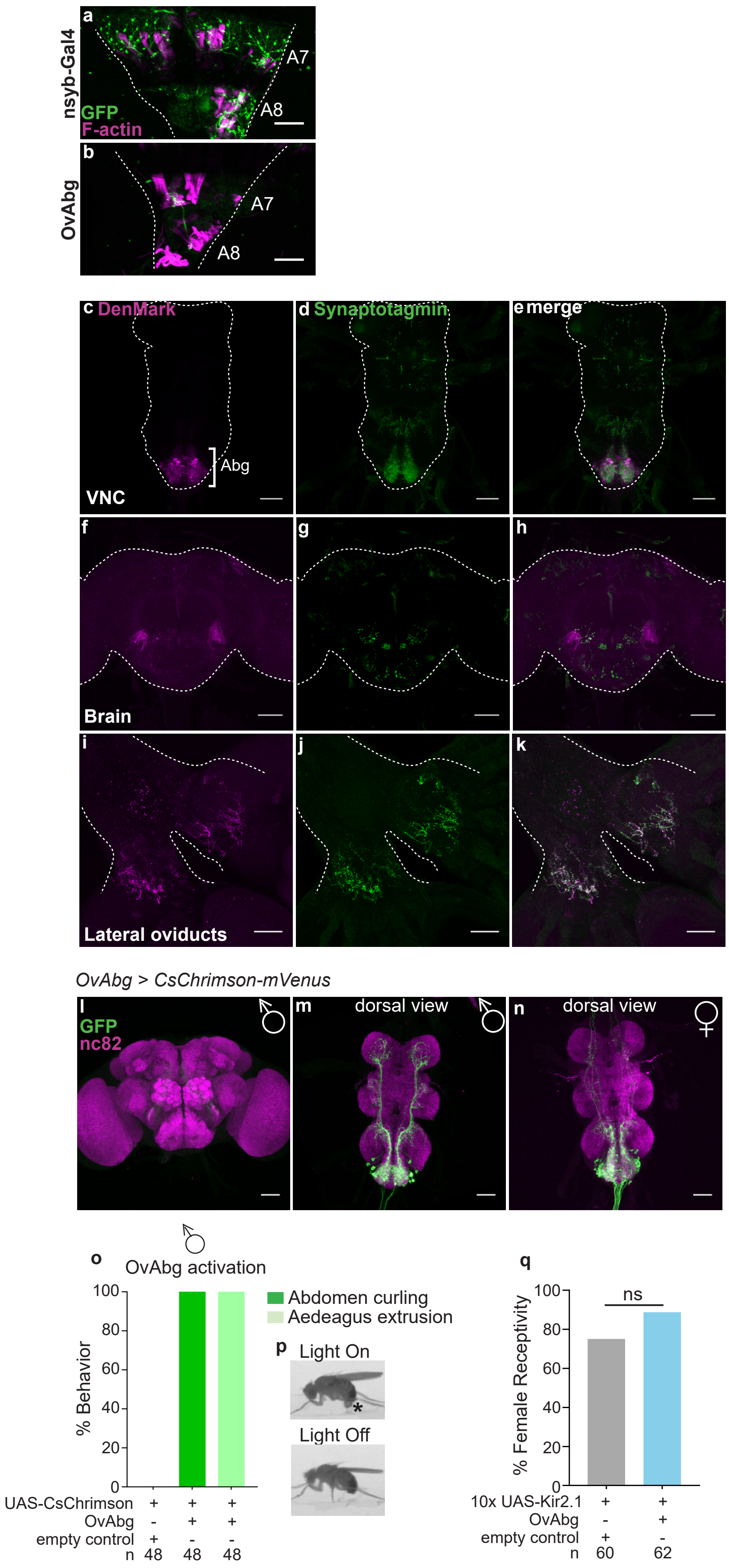


**Supplementary Fig. 1.**

**a** and **b** Confocal images of female abdominal muscles in the Nsyb-Gal4 (**a**) and OvAbg line (**b**). Neuronal innervations are stained with anti-GFP (green) and muscle fibers with anti-F-actin (magenta).

A7 and A8 indicate the position of the abdominal segments. Nsyb-Gal4 line expression is shown for comparison with the OvAbg line expression. Anti-GFP is targeting the fluorescent protein GFP and Venus from Nsyb-Gal4 > mCD8::GFP and OvAbg > CsChrimson-mVenus flies. Scale bars a) and b) 50 μm. **c-k** Confocal images of OvAbg neuronal polarity in the female VNC (**c-e**), brain (**f-h**) and reproductive system (**i-k**). Dendrites (inputs) are labelled using the somatodendritic marker, DenMark, and axons (outputs) are labelled using the synaptic vesicle marker, Synaptotagmin. Anti-GFP is targeting EGFP-tagged Synaptotagmin and anti-DsRed is targeting mCherry-tagged DenMark. Scale bars c-k) 50 μm. **l-n** Confocal images of male brain (**l**) as well as male (**m**) and female (**n**) dorsal view of VNC showing OvAbg neurons and corresponding innervations stained with anti-GFP (green) to reveal the anatomy and nc82 for synapses. Anti-GFP is targeting the fluorescent protein Venus from OvAbg > CsChrimson-mVenus expressing flies. Scale bars l), m) and n) 50 μm. **o** Percentage of stimulation events in which male OvAbg flies displayed abdomen curling and Aedeagus extrusion behaviours during photoactivation with CsChrimson. n = 48 (control) and 48 (OvAbg) stimulations. **p** Video snapshot (lateral view) of OvAbg male displaying abdomen curling and Aedeagus extrusion (asterisk) behaviours in response to the stimulation with CsChrimson (top). A snapshot of the same male during a light off period (below) is also shown for comparison. **q** Percentage of receptive females during inhibition of OvAbg neurons. n = 60 (control) and 62 (OvAbg) females. Fisher´s exact test, ns $p \geq 0.05$.

Supplementary Fig. 2

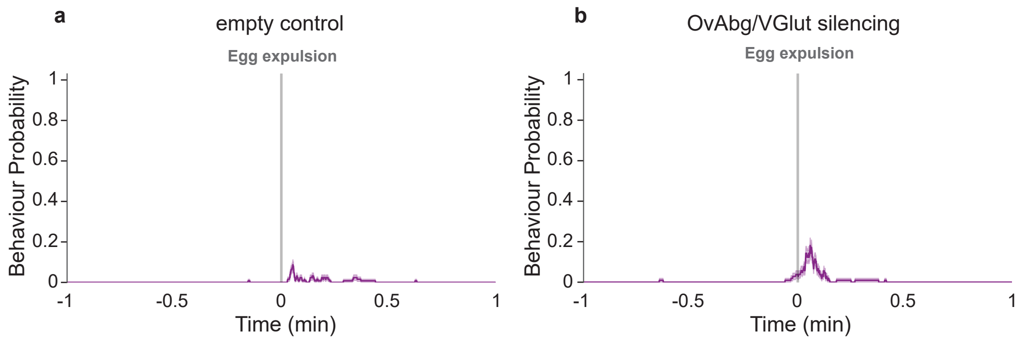


**Supplementary Fig. 2**

**a** and **b** Probabilities of grooming terminalia behaviour during a 1-min time window around egg expulsion for (**a**) control and (**b**) OvAbg/VGlut silenced flies. Time = 0 minutes marks the moment of egg expulsion (represented by the grey vertical line). n = 126 (control) and n = 105 (OvAbg/VGlut) egg expulsions.

#### Supplementary Table 1. Fly stocks

| Stocks | Source | Reference |
|---|---|---|
| *Canton S (CS)* | Lab stock | - |
| *w-; empty-p65-AD(attp40); +* | Bloomington Drosophila Stock Center (BDSC), 71210 | Hampel et al[1] |
| *w-; VT038154-p65.AD(attp40); +* | BDSC, 74056 | Tirian et al[2] |
| *w-;+; $dsx^{DBD}$* | - | Pavlou et al[3] |
| *w-; +; R57C10-GAL4(attp2)* | BDSC, 39171 | Jenett et al[4] |
| *$y^1$w-; Gad1(MI09277)-LexA:QFAD; +* | BDSC, 60324 | Diao et al[5] |
| *$y^1$w-; ChaT(MI04508)-LexA:QFAD; +* | BDSC, 60319 | Diao et al[5] |
| *$y^1$w-; VGlut(MI04979)-LexA:QFAD; +* | BDSC, 60314 | Diao et al[5] |
| *w-; +; 20xUAS-CsChrimson-mVenus(attp2)* | BDSC, 55136 | Klapoetke et al[6] |
| *w-, 20xUAS-CsChrimson-mVenus(attP18);+;+* | BDSC, 55134 | Markstein et al[7] |
| *w-;; 20xUAS>Stop>CsChrimson-mVenus(attp2)* | FLP-out version provided by Vivek Jayaraman | Klapoetke et al[6] |
| *DL; +; 10xUAS-Kir2.1-EGFP* | Provided by Moita laboratory | - |
| *w-;UAS>Stop>Kir2.1-EGFP(VIE-19A);+* | - | Asahina et al[8] |
| *w-; +; 20xUAS-GtACR1(attp2)* | - | Mohammad et al[9] |
| *w-; 8xLexAop2-FLP(attp40); +* | BDSC, 55820 | Pan et al[10] |
| *w-; +; 8xLexAop2-FLP(attp2)* | BDSC, 55819 | Pan et al[10] |
| *w-; +; 20xUAS-mCD8::GFP(attp2)* | BDSC, 32194 | Pfeiffer at al[11] |
| *w-; UAS-DenMark2, UAS-syt.EGFP* | BDSC, 33064 | Nicolai et al[12] |

### Supplementary Table 2. Full genotypes of flies used in experiments

| Figures | Genotypes |
|---|---|
| Figure 1 | Canton S (CS) |
| Figure 2a-j | Control: w-; empty-p65-AD(attp40) / +; $dsx^{DBD}$ / 20xUAS-CsChrimson-mVenus(attp2)<br>Test: w-; VT038154-p65-AD(attp40) / +; $dsx^{DBD}$ / 20xUAS-CsChrimson-mVenus(attp2) |
| Figure 2k-m | Control: w- / DL; empty-p65-AD(attp40) / +; $dsx^{DBD}$ / 10xUAS-Kir2.1-EGFP<br>Test: w- / DL; VT038154-p65-AD(attp40) / +; $dsx^{DBD}$ / 10xUAS-Kir2.1-EGFP |
| Figure 3 | Control: w-; empty-p65-AD(attp40) / +; $dsx^{DBD}$ / 20xUAS-GtACR1(attp2)<br>Test: w-; VT038154-p65-AD(attp40) / +; $dsx^{DBD}$ / 20xUAS-GtACR1(attp2) |
| Figure 4a-c, e-g | Control: w-; empty-p65-AD(attp40) / 8xLexaop2-FLP; $dsx^{DBD}$, UAS>Stop>CsChrimson-mVenus(attp2) / Gad1- LexA:QFAD (MI09277)<br>Test: w-; VT038154-p65-AD(attp40) / 8xLexaop2-FLP; $dsx^{DBD}$, UAS>Stop>CsChrimson-mVenus(attp2) / Gad1- LexA:QFAD (MI09277) |
| Figure 4d | Control: w-; empty-p65-AD(attp40), UAS>Stop>Kir2.1-EGFP(VIE-19A) / 8xLexaop2-FLP(attp40); $dsx^{DBD}$ / Gad1- LexA:QFAD (MI09277)<br>Test: w-; VT038154-p65-AD(attp40), UAS>Stop>Kir2.1-EGFP(VIE-19A) / 8xLexaop2-FLP(attp40); $dsx^{DBD}$ / Gad1- LexA:QFAD (MI09277) |
| Figure 5a-c, f-i | Control: w-; empty-p65-AD(attp40) / 8xLexaop2-FLP(attp40); $dsx^{DBD}$, UAS>Stop>CsChrimson-mVenus(attp2) / ChaT-LexA:QFAD (MI04508)<br>Test: w-; VT038154-p65-AD(attp40) / 8xLexaop2-FLP(attp40); $dsx^{DBD}$, UAS>Stop>CsChrimson-mVenus(attp2) / ChaT-LexA:QFAD (MI04508) |
| Figure 5d-e | Control: w-; empty-p65-AD(attp40), UAS>Stop>Kir2.1-EGFP(VIE-19A) / 8xLexaop2-FLP(attp40); $dsx^{DBD}$ / ChaT-LexA:QFAD (MI04508)<br>Test: w-; VT038154-p65-AD(attp40), UAS>Stop>Kir2.1-EGFP(VIE-19A) / 8xLexaop2-FLP(attp40); $dsx^{DBD}$ / ChaT-LexA:QFAD (MI04508) |
| Figure 5j | 1. w-; VT038154-p65-AD(attp40) / +; $dsx^{DBD}$ / 20xUAS-CsChrimson-mVenus(attp2)<br>2. w-; VT038154-p65-AD(attp40) / 8xLexaop2-FLP(attp40); $dsx^{DBD}$, UAS>Stop>CsChrimson-mVenus(attp2) / ChaT-LexA:QFAD (MI04508) |
| Figure 6a-e | Control: w-; empty-p65-AD(attp40) / VGlut-LexA:QFAD (MI04979); $dsx^{DBD}$, UAS>Stop>CsChrimson-mVenus(attp2) / 8xLexaop2-FLP(attp2)<br>Test: w-; VT038154-p65-AD(attp40) / VGlut-LexA:QFAD (MI04979); $dsx^{DBD}$, UAS>Stop>CsChrimson-mVenus(attp2) / 8xLexaop2-FLP(attp2) |
| Figure 6f-p | Control: w-; empty-p65-AD(attp40), UAS>Stop>Kir2.1-EGFP(VIE-19A) / VGlut-LexA:QFAD (MI04979); $dsx^{DBD}$ / 8xLexaop2-FLP(attp2)<br>Test: w-; VT038154-p65-AD(attp40), UAS>Stop>Kir2.1-EGFP(VIE-19A) / VGlut-LexA:QFAD (MI04979); $dsx^{DBD}$ / 8xLexaop2-FLP(attp2) |
| Supplementary Figure 1a-b | a. w-; +; GMR57C10-GAL4(attp2) / 20xUAS-mCD8::GFP(attp2) (nsyb)<br>b. w-; VT038154-p65-AD(attp40) / +; $dsx^{DBD}$ / 20xUAS-CsChrimson-mVenus(attp2) |
| Supplementary Figure 1c-k | w-; VT038154-p65-AD(attp40) / UAS-Synaptotagmin-EGFP, UAS-DenMark2-mCherry; $dsx^{DBD}$ / + |
| Supplementary Figure 1l-n | w-, 20xUAS-CsChrimson-mVenus(attP18); VT038154-p65-AD(attp40) / +; $dsx^{DBD}$ / + |
| Supplementary Figure 1o-p | Control: w-; empty-p65-AD(attp40) / +; $dsx^{DBD}$ / 20xUAS-CsChrimson-mVenus(attp2)<br>Test: w-; VT038154-p65-AD(attp40) / +; $dsx^{DBD}$ / 20xUAS-CsChrimson-mVenus(attp2) |
| Supplementary Figure 1q | Control: w- / DL; empty-p65-AD(attp40) / +; $dsx^{DBD}$ / 10xUAS-Kir2.1-EGFP<br>Test: w- / DL; VT038154-p65-AD(attp40) / +; $dsx^{DBD}$ / 10xUAS-Kir2.1-EGFP |
| Supplementary Figure 2a-b | Control: w-; empty-p65-AD(attp40), UAS>Stop>Kir2.1-EGFP(VIE-19A) / VGlut-LexA:QFAD (MI04979); $dsx^{DBD}$ / 8xLexaop2-FLP(attp2)<br>Test: w-; VT038154-p65-AD(attp40), UAS>Stop>Kir2.1-EGFP(VIE-19A) / VGlut-LexA:QFAD (MI04979); $dsx^{DBD}$ / 8xLexaop2-FLP(attp2) |

### Supplementary Table 3. Statistical details related to main and supplementary figures

| Figure | Groups | N | Normally distributed* | Equal variance** | Statistical test | p-value |
|---|---|---|---|---|---|---|
| 2j | a) OvAbg Kir2.1 silencing<br>b) empty control | 45<br>55 | no<br>no | no | Mann-Whitney test | < 0.0001 (****) |
| 2k | a) OvAbg Kir2.1 silencing<br>b) empty control | 32<br>36 | NA<br>NA | NA<br>NA | Fisher´s exact test | < 0.0001 (****) |
| 3b | Pre: a) OvAbg GtACR1 silencing<br>b) empty control | 19<br>8 | no<br>yes | yes | Mann-Whitney test | = 0.1340 (ns) |
| | Silencing: c) OvAbg GtACR1 silencing<br>d) empty control | 19<br>8 | no<br>yes | no | Mann-Whitney test | = 6.44E-06 (****) |
| | Post: e) OvAbg GtACR1 silencing<br>f) empty control | 19<br>8 | no<br>yes | yes | Mann-Whitney test | = 0.0151 (*) |
| 3c | Pre: a) OvAbg GtACR1 silencing<br>b) empty control | 19<br>8 | no<br>yes | yes | Mann-Whitney test | = 0.1341 (ns) |
| | Silencing: c) OvAbg GtACR1 silencing<br>d) empty control | 19<br>8 | no<br>yes | yes | Mann-Whitney test | = 8.11E-06 (****) |
| | Post: e) OvAbg GtACR1 silencing<br>f) empty control | 19<br>8 | no<br>yes | yes | Mann-Whitney test | = 0.0106 (*) |
| 3d | Pre: a) OvAbg GtACR1 silencing<br>b) empty control | 19<br>8 | yes<br>yes | yes | t-test | = 0.2259 (ns) |
| | Silencing: c) OvAbg GtACR1 silencing<br>d) empty control | 19<br>8 | yes<br>yes | no | t-test with Welch´s correction | = 0.0023 (**) |
| | Post: e) OvAbg GtACR1 silencing<br>f) empty control | 19<br>8 | no<br>yes | yes | Mann-Whitney test | = 0.0106 (*) |
| 3e | Pre: a) OvAbg GtACR1 silencing<br>b) empty control | 19<br>8 | no<br>yes | yes | Mann-Whitney test | = 0.1341 (ns) |
| | Silencing: c) OvAbg GtACR1 silencing<br>d) empty control | 19<br>8 | yes<br>yes | no | t-test with Welch´s correction | = 0.0023 (**) |
| | Post: e) OvAbg GtACR1 silencing<br>f) empty control | 19<br>8 | no<br>yes | yes | Mann-Whitney test | = 0.0106 (*) |
| 3f | Pre: a) OvAbg GtACR1 silencing<br>b) empty control | 19<br>8 | yes<br>yes | yes | t-test | = 0.0680 (ns) |
| | Silencing: c) OvAbg GtACR1 silencing<br>d) empty control | 19<br>8 | yes<br>yes | no | t-test with Welch´s correction | = 0.0013 (**) |
| | Post: e) OvAbg GtACR1 silencing<br>f) empty control | 19<br>8 | no<br>yes | yes | Mann-Whitney test | = 0.0151 (*) |
| 3g | Pre: a) OvAbg GtACR1 silencing<br>b) empty control | 19<br>8 | yes<br>yes | yes | t-test | = 0.9607 (ns) |
| | Silencing: c) OvAbg GtACR1 silencing<br>d) empty control | 19<br>8 | no<br>yes | yes | Mann-Whitney test | = 0.4893 (ns) |
| | Post: e) OvAbg GtACR1 silencing<br>f) empty control | 19<br>8 | no<br>yes | yes | Mann-Whitney test | = 0.0286 (*) |
| 3h | Pre: a) OvAbg GtACR1 silencing<br>b) empty control | 15<br>7 | no<br>yes | yes | Mann-Whitney test | = 0.0604 (ns) |
| | Silencing: c) OvAbg GtACR1 silencing<br>d) empty control | 19<br>8 | yes<br>yes | no | t-test with Welch´s correction | = 9.17E-05 (****) |
| | Post: e) OvAbg GtACR1 silencing<br>f) empty control | 7<br>6 | no<br>no | yes | Mann-Whitney test | = 0.0058 (**) |
| 3i | Pre: a) OvAbg GtACR1 silencing<br>b) empty control | 19<br>8 | yes<br>yes | yes | t-test | = 0.6449 (ns) |
| | Silencing: c) OvAbg GtACR1 silencing<br>d) empty control | 19<br>8 | no<br>yes | yes | Mann-Whitney test | = 0.0440 (*) |
| | Post: e) OvAbg GtACR1 silencing<br>f) empty control | 19<br>8 | no<br>yes | yes | Mann-Whitney test | = 0.4784 (ns) |
| 3j | Pre: a) OvAbg GtACR1 silencing<br>b) empty control | 19<br>8 | no<br>yes | yes | Mann-Whitney test | = 0.3355 (ns) |
| | Silencing: c) OvAbg GtACR1 silencing<br>d) empty control | 14<br>8 | no<br>yes | yes | Mann-Whitney test | = 0.0220 (*) |
| | Post: e) OvAbg GtACR1 silencing<br>f) empty control | 16<br>7 | yes<br>yes | yes | t-test | = 0.4843 (ns) |
| 3l | Pre: a) OvAbg GtACR1 silencing<br>b) empty control | 19<br>8 | yes<br>yes | yes | t-test | = 0.8649 (ns) |
| | Silencing: c) OvAbg GtACR1 silencing<br>d) empty control | 19<br>8 | no<br>yes | yes | Mann-Whitney test | = 0.0002 (***) |
| | Post: e) OvAbg GtACR1 silencing<br>f) empty control | 19<br>8 | no<br>no | yes | Mann-Whitney test | = 0.2848 (ns) |
| 3m | Pre: a) OvAbg GtACR1 silencing<br>b) empty control | 19<br>8 | no<br>no | yes | Mann-Whitney test | = 0.1862 (ns) |
| | Silencing: c) OvAbg GtACR1 silencing<br>d) empty control | 19<br>8 | yes<br>no | no | Mann-Whitney test | = 0.0001 (****) |
| | Post: e) OvAbg GtACR1 silencing<br>f) empty control | 19<br>8 | no<br>no | yes | Mann-Whitney test | = 0.2069 (ns) |
| 3n | Pre: a) OvAbg GtACR1 silencing<br>b) empty control | 19<br>8 | no<br>yes | yes | Mann-Whitney test | = 0.0872 (ns) |
| | Silencing: c) OvAbg GtACR1 silencing<br>d) empty control | 19<br>8 | no<br>yes | yes | Mann-Whitney test | = 0.0084 (**) |
| | Post: e) OvAbg GtACR1 silencing<br>f) empty control | 19<br>8 | no<br>no | yes | Mann-Whitney test | = 0.0887 (ns) |
| 4d | a) OvAbg/Gad1 Stop>Kir2.1 silencing<br>b) empty control | 49<br>46 | yes<br>no | no | Mann-Whitney test | = 0.9276 (ns) |
| 4e | a) OvAbg/Gad1 Stop>Chrimson activation<br>b) empty control | 40<br>29 | no<br>no | no | Mann-Whitney test | < 0.0001 (****) |
| 4f | a) OvAbg/Gad1 Stop>Chrimson activation<br>b) empty control | 35<br>25 | NA<br>NA | NA<br>NA | Fisher´s exact test | < 0.0001 (****) |
| 5d | a) OvAbg/Cha Stop>Kir2.1 silencing<br>b) empty control | 25<br>45 | no<br>yes | no | Mann-Whitney test | < 0.0001 (****) |
| 5e | a) OvAbg/Cha Stop>Kir2.1 silencing<br>b) empty control | 19<br>28 | NA<br>NA | NA<br>NA | Fisher´s exact test | < 0.0001 (****) |
| 5h | a) OvAbg/Cha Stop>Chrimson activation<br>b) empty control | 24<br>25 | NA<br>NA | NA<br>NA | Fisher´s exact test | < 0.0001 (****) |
| 5j | a) OvAbg/Cha Stop>Chrimson activation<br>b) OvAbg Chrimson activation | 24<br>9 | no<br>yes | yes | Mann-Whitney test | < 0.0001 (****) |

| | | | | | | |
|---|---|---|---|---|---|---|
| 6d | *a)* OvAbg/VGlut Stop>Chrimson activation (Mated)<br>b) OvAbg/VGlut Stop>Chrimson activation (Virgin) | 138<br>79 | NA<br>NA | NA<br>NA | Fisher´s exact test | < 0.0001 (****) |
| 6f | a) OvAbg/VGlut Stop>Kir2.1 silencing<br>b) empty control | 38<br>26 | no<br>yes | yes | Mann-Whitney test | = 0.0003 (***) |
| 6g | a) OvAbg/VGlut >Kir2.1 silencing<br>b) empty control | 38<br>26 | no<br>yes | no | Mann-Whitney test | = 0.0002 (***) |
| 6h | a) OvAbg/VGlut >Kir2.1 silencing<br>b) empty control | 38<br>26 | no<br>no | no | Mann-Whitney test | = 4.13E-06 (****) |
| 6i | a) OvAbg/VGlut >Kir2.1 silencing<br>b) empty control | 38<br>26 | no<br>no | no | Mann-Whitney test | = 2.99E-06 (****) |
| 6j | a) OvAbg/VGlut >Kir2.1 silencing<br>b) empty control | 38<br>26 | no<br>yes | yes | Mann-Whitney test | =0.2132 (ns) |
| 6k | a) OvAbg/VGlut >Kir2.1 silencing<br>b) empty control | 38<br>26 | no<br>no | no | Mann-Whitney test | = 0.0124 (*) |
| 6l | a) OvAbg/VGlut >Kir2.1 silencing<br>b) empty control | 38<br>26 | no<br>no | yes | Mann-Whitney test | = 0.2918 (ns) |
| 6m | a) OvAbg/VGlut >Kir2.1 silencing<br>b) empty control | 38<br>26 | no<br>no | yes | Mann-Whitney test | = 0.3962 (ns) |
| 6n | a) OvAbg/VGlut >Kir2.1 silencing<br>b) empty control | 38<br>26 | no<br>no | yes | Mann-Whitney test | = 0.0693 (ns) |
| 6o | a) OvAbg/VGlut >Kir2.1 silencing<br>b) empty control | 38<br>26 | no<br>yes | no | Mann-Whitney test | = 0.4001 (ns) |
| 6p | a) OvAbg/VGlut >Kir2.1 silencing<br>b) empty control | 38<br>26 | no<br>no | yes | Mann-Whitney test | = 0.0005 (***) |
| S1q | a) OvAbg Kir2.1 silencing<br>b) empty control | 62<br>60 | NA<br>NA | NA<br>NA | Fisher´s exact test | 0.0608 (ns) |

NA = not applicable

* To test for normal distribution: Shapiro's Test, D'Agostino's Test, Anderson-Darling test and Kolmogorov-Smirnov test.

** To test for the homogeneity of variance: Levene´s test for non-normally distributed samples and Bartlett´s test for normally distributed samples.
